# Serial Dependence Predicts Generalization in Perceptual Learning

**DOI:** 10.1101/2025.02.06.636846

**Authors:** Noga Pinchuk-Yacobi, Dov Sagi, Yoram S. Bonneh

## Abstract

Visual perception is shaped by recent experience, but how these momentary influences accumulate to support long-term learning and generalization remains unclear. Here, we asked whether short-term memory traces, attractive serial-dependence effects (SDEs), promote learning generalization. We re-analyzed over 200,000 trials from observers trained on a visual texture-discrimination task under three conditions that differentially modulated generalization. Under certain conditions, SDEs reached further back in time than previously reported and persisted after eight days of practice, despite the non-informative nature of past stimuli. Observers in conditions that promoted generalization displayed larger long-range SDEs, and individual SDE magnitude predicted transfer of learning across locations. We propose that SDE is associated with learning flexibility, providing a principled framework for when and why perceptual learning generalizes, which is central to theories of cognitive flexibility. Attractive serial dependence is not an extra mechanism in this model—it is the behavioral footprint of ongoing template plasticity required for flexibility in changing environments.

---

Perceptual learning (PL) and serial dependence effects (SDEs) are two fundamental processes that shape how we perceive and interpret sensory information. Although both rely on perceptual memory traces, they operate over distinct timescales and have been traditionally studied separately. PL leads to long-lasting improvements in sensory discrimination following repeated training (Sagi, 2011). However, in many cases, improvements remain limited to the trained stimulus or location, while in others, learning generalizes to new contexts. What determines whether learning stays local or generalizes is still not fully understood (Cheng et al., 2025; Lu & Dosher, 2022).

SDEs reflect short-term biases that pull current perceptual judgments toward what was recently seen or chosen (Abrahamyan et al., 2016; Braun et al., 2018; Falmagne et al., 1975; Fischer & Whitney, 2014; Fornaciai & Park, 2018; Liberman et al., 2014; Suárez-Pinilla et al., 2018; Urai et al., 2019). These biases are often interpreted as reflecting a Bayesian integration process that combines prior information with current input to enhance perceptual stability and efficiency, particularly under uncertainty, based on the assumption that the natural environment is typically stable (Cicchini et al., 2018; Fischer & Whitney, 2014; Manassi & Whitney, 2024). However, in tasks where stimuli vary randomly across trials, such biases can actually impair performance, yet they persist. This suggests they may serve an additional role beyond what was previously proposed. One possibility is that they reflect ongoing trial-by-trial updates to internal decision templates that persist over time and may bridge short-term memory and long-term learning (Fritsche et al., 2017; Pascucci et al., 2019).

In this work, we examine how serial dependence interacts with perceptual learning using the texture discrimination task (TDT; Karni & Sagi, 1991). The role of serial dependence in the TDT is likely complex, potentially influencing both immediate perceptual judgments and long-term learning dynamics. In the short term, biases toward the orientation of prior targets, randomly varying in typical experimental setups, and thus irrelevant to the current trial, may impair performance by introducing decision noise. At the same time, temporal integration of stimulus representations across trials may help reduce uncertainty and extract structure from noisy input. Normative analyses predict that the benefit of engaging costlier, memory-dependent integration should peak at intermediate uncertainty, while being limited in highly certain conditions (Tavoni et al., 2022). Over extended training, serial dependence may evolve within and across daily sessions as learning progresses. Serial dependence may also arise as a result of trial by trial network updates during learning (Petrov et al., 2006, 2005), depending on how the effect of recent stimuli is weighted during network updates. Specifically, we ask whether serial dependence can serve as a marker of short-term temporal integration that contributes to long-term generalization of learning. This idea builds on the hypothesis that broader temporal integration, covering a larger network state space, may reduce overfitting to local stimulus features and allow learning to generalize more broadly.

To test this, we reanalyzed data from a large-scale perceptual learning study in which observers practiced the TDT under three training conditions designed to modulate the degree of learning generalization (Harris et al., 2012). The study showed that learning generalized to a new, untrained location when targets appeared randomly across two locations or were intermixed with target-less (dummy’) trials. In contrast, learning became location-specific when targets consistently appeared in a single location in all trials, likely due to increased sensory adaptation (Karni & Sagi, 1991). These findings were attributed to differences in adaptation state: unadapted networks supported spatial transfer, whereas adaptation induced localized plasticity that constrained generalization. However, the memory mechanisms underlying this flexibility remain unclear. By examining serial dependence within this paradigm, we aim to gain further insight into these mechanisms and better understand how recent visual experiences influence both immediate perceptual reports and long-term learning outcomes. Specifically, we ask whether training conditions that promote generalization are associated with stronger or longer-lasting SDEs, suggesting a new role for serial dependence in contributing to broader learning transfer. We consider a learning mechanism where adaptation dependent inhibition controls decision template updates, and by that learning flexibility. On this account, serial dependence is a consequence of network plasticity.

## Methods

We reanalyzed data from 50 observers who participated in the texture discrimination task (TDT) as described by Harris Gliksberg, and Sagi (2012). In this dual-task experiment, observers identified the orientation (vertical or horizontal) of a target composed of three peripheral diagonal lines embedded within a uniform background of horizontal lines while simultaneously performing a forced-choice letter discrimination task (T vs L at the center of the stimulus) to maintain fixation (Figure 1A). The experiment consisted of four daily sessions with the target presented at one location (or two fixed locations in the 2loc condition), followed by four additional daily sessions at a second location (or a second pair of fixed locations in the 2loc condition). Performance on the TDT task, the TDT threshold, is quantified as the SOA (see Figure 1) that yields 78% correct discrimination (SOA_threshold_). The reaction time (RT) used in the analysis was defined as RT(TDT) − RT(fixation task), where RT for each task was measured from stimulus onset.

**Figure 1.**
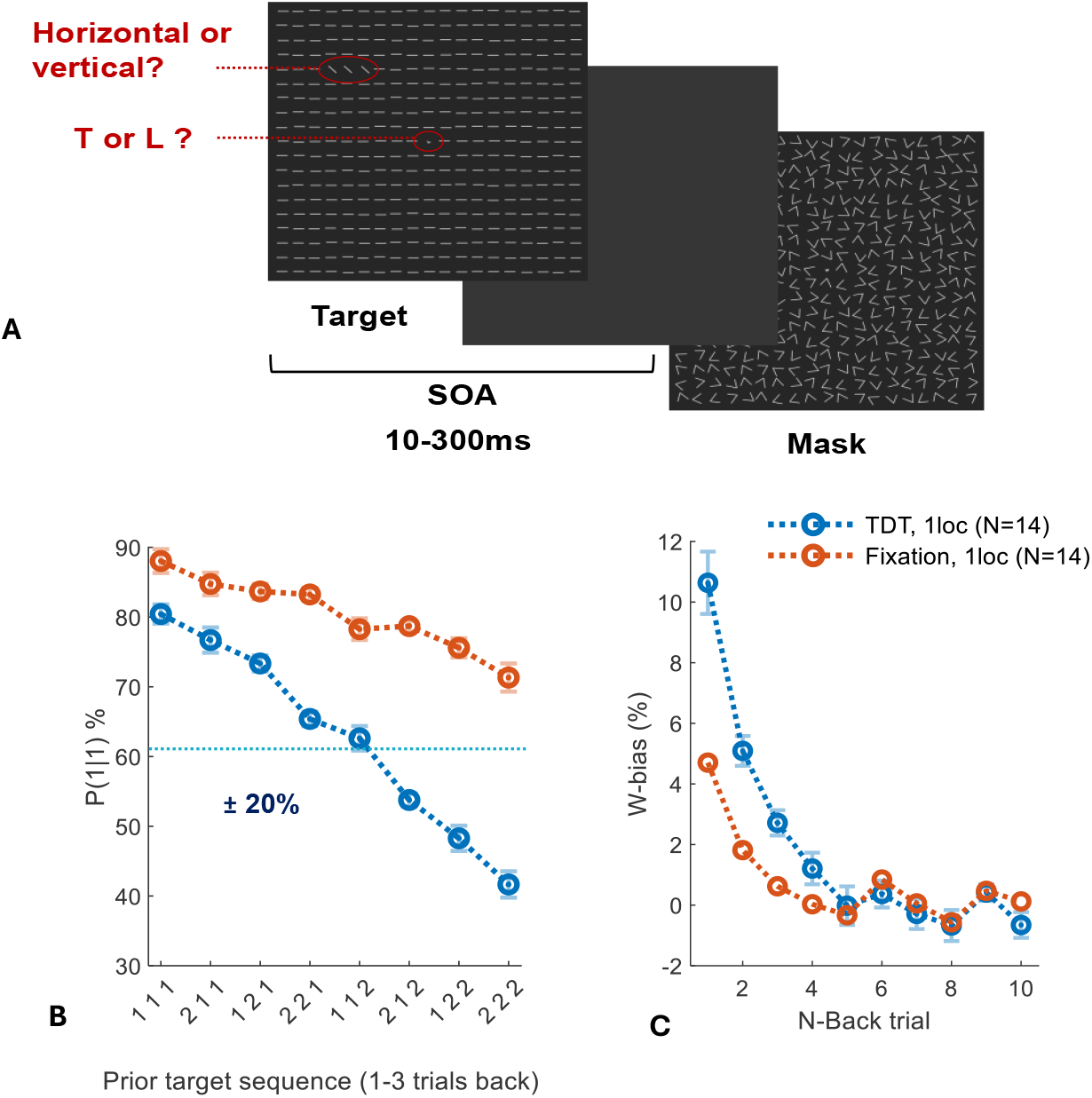
Texture Discrimination Task (TDT) and Serial Dependence Effect (SDE) (A) TDT Trial Sequence: Observers identified the orientation (vertical or horizontal) of a target composed of three diagonal lines embedded within a background of horizontal lines while simultaneously performing a forced-choice letter discrimination task (T vs L) to maintain fixation. The 10 ms target frame was followed by a 100 ms patterned mask, with the stimulus onset asynchrony (SOA) between target and mask (10 to 300 ms) randomized across trials. (B) SDE via history sequence: Correct identifications (‘Hits’) increased by approximately 20% when the current target orientation matched the preceding three orientations (e.g., ‘1’ preceded by ‘111’) and decreased by about 20% with mismatching orientations (e.g., ‘1’ preceded by ‘222’), relative to average performance (dotted light blue line). ‘1’ and ‘2’ represent the two possible orientations, either vertical and horizontal, or vice versa. The fixation task (red) showed much smaller biases, likely due to its higher overall performance. (C) SDE via Linear Mixed Effects (LME) Weights (W-bias, %): Influence of 1–10 back trials on current report. Summing the W-bias values (%) from the 1st, 2nd, and 3rd prior trials corresponds to the ±20% bias for 1–3 back trials shown in panel B. Panels B and C include data from the 1loc condition (N=14) pooled across all training days (Days 1–8; 4 days at the first location and 4 days at the second location to assess generalization), including only trials with low-visibility current targets (SOA < SOA_threshold_ + 20 ms, calculated on a per-subject basis). Blue represents the texture discrimination task, and red indicates the letter discrimination (fixation control) task.

Participants were assigned to one of three experimental conditions:

- **1loc condition**: The target consistently appeared at the same fixed location across all trials.
- **2loc condition**: The target randomly appeared at one of two diagonally opposite locations with equal eccentricity, alternating between trials.
- **Dummy condition**: The target appeared at a fixed location, but genuine target trials were randomly interleaved with ‘dummy’ trials, where no target was present (replaced by background elements).

For detailed specifications of the groups assigned to each condition, see Supplementary Table S1.

We quantified serial dependence effects (SDE) to examine how prior visual experiences influence current perceptual reports (Figure 1B), and whether these dependencies affect learning specificity across experimental conditions. To estimate SDE, we fit a linear mixed-effects (LME) model (Figure 1C) that evaluated the influence of prior target stimuli (up to’ 10 trials back) on current reports. For each lag n, the model estimates a coefficient *W*_*n*_; we refer to this coefficient simply as W-bias. These coefficients represent the magnitude of the serial-dependence effect for each n-back stimulus. We modeled the report probability as

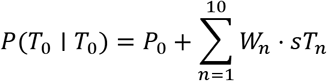

Where:

- *P*(*T*_0_ | *T*_0_): probability of reporting target *T*_0_ ∈ {+1, −1} when the target is *T*_0_.
- *P*_0_: history-independent baseline report probability.
- *sT*_*n*_: +1 if the *n*-back target’s orientation matches the current target (e.g., both vertical or both horizontal), −1 otherwise.
- *W*_*n*_(W-bias): change in report probability (bias) associated with the *n*-back target.

Fixed effects accounted for the orientations of these stimuli, while a random intercept captured individual differences across observers. Interaction terms were excluded, as statistically significant interactions were small and inconsistent. To preserve the precise temporal structure of the data, all trials were included in the sequential n-back count across all experimental conditions. In the Linear Mixed Effects (LME) analysis, we modeled these trial types using distinct regressors: each n-back lag included separate predictors for visible and invisible targets, further differentiated by trial type (dummy vs. target) and relative location (ipsilateral vs. contralateral) where applicable. The SDE values reported here reflect only the influence of relevant target-present history trials; the effects of other history types (e.g., dummy trials), while estimated to ensure the temporal integrity of the model, are not presented.

To systematically quantify serial dependence across different temporal scales, we defined three summary measures:

- **SDE-all:** The cumulative bias from the 10 preceding trials (1–10 back), capturing the total influence of recent history on current perception.
- **SDE-recent:** The cumulative bias from trials 1–3 back, reflecting the effect of very recent stimuli.
- **SDE-distant:** The cumulative bias from trials 4–6 back, representing the influence of more distant past trials.

All measures are expressed as percentage changes in report probability. For most analyses and figures, we filtered trials to include those that produced stronger SDEs. The specific filtering applied is motivated by technical issues. When using percent correct as a measure of performance, bias cannot be reliably estimated at or near ceiling performance, as correct responses leave little room for bias to manifest. However, easy’ trials can be taken as reliable references to condition the bias on, since low performance introduces uncertainty as for the perceptual effect of the reference orientation. Specifically, trials in which current targets were barely visible (SOA < SOA_threshold_ + 20 ms), prior targets were highly visible (SOA > SOA_threshold_), and both appeared at the same location across trials (Figure 2). The exception was the SDE derived from the history sequence analysis (Figure 1B), where prior-target visibility could not be filtered. Therefore, when comparing this with the LME-based estimate (Figure 1C), only the current-target visibility filter was applied. Finally, to verify that our findings are not limited to these filtering choices, we also conducted control analyses including all prior-trial history regardless of visibility; these results are presented in Supplementary Figure S3 and confirm the robustness of our main findings.

**Figure 2.**
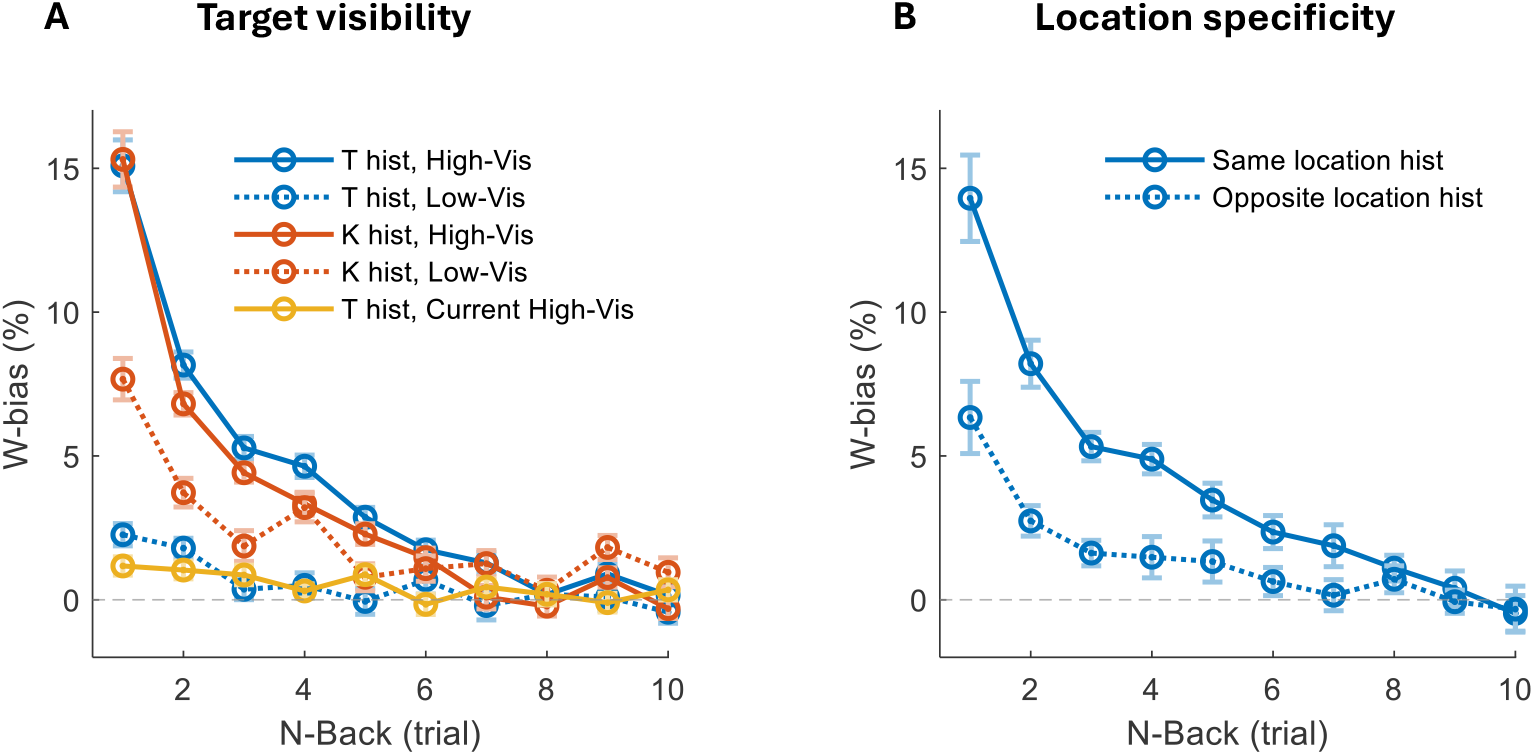
Serial dependence effects across trial history. (A) Target visibility: SDEs (W-bias) were strongest when current targets had low visibility (high uncertainty) and prior targets had high visibility. Blue lines show target (T) histories and red lines show key (K) histories, with solid vs. dotted lines indicating prior high-vs. low-visibility trials. Visibility had a much weaker effect on key histories compared to target histories, with key-driven SDEs remaining high even when prior targets were invisible. The yellow line shows target history with high visibility in both history and current trials, demonstrating that SDEs are strongly reduced when current targets are highly visible (certainty) (*N = 50*). (B) Location specificity: SDEs were larger when history originated from the same location as the current target compared to a diagonally opposite location (2loc condition, *N = 14*). Error bars represent the standard error of the mean. The horizontal dashed line indicates zero bias.

For individual-level analyses assessing the relationship between SDE strength and TDT learning and generalization, we accounted for individual differences in SOA_threshold_ by equalizing the number of highly visible prior trials and restricting trials to SOAs between SOA_threshold_ and SOA_threshold_ + 140 ms.

Because filtering on highly visible prior targets means observers usually reported them correctly, the key-history sequence closely tracked the target-history sequence, producing highly similar SDEs (Figure 2A, high-vis history). Accordingly, we present only Target-history results and omit Key-history plots for brevity.

For computing correlations on individual participants (Figures 4, 7), we used orthogonal-regression without excluding outliers, then, we used the Pearson correlation to compute R and p-value. Confidence sleeves around regression lines were estimated using nonparametric bootstrap resampling (1,000 iterations). For each bootstrap sample, an orthogonal regression line was fit, and the 95% confidence sleeve was defined by the 2.5th and 97.5th percentiles of the predicted values across resamples. For non-significant correlations, we omitted the orthogonal regression line but kept the confidence sleeves.

Data Availability. The data that support the findings of this study are available from the corresponding author upon request.

Ethics. This manuscript presents a secondary analysis of previously published human behavioural data from Harris, Gliksberg & Sagi (2012). Original procedures received ethics approval, were conducted in accordance with the Declaration of Helsinki, and all participants provided written informed consent.

## Results

### Serial Dependence Effects (SDEs)

Observers’ reports showed a significant 15% bias toward the orientation of the immediately preceding target (1-back; W1), indicating serial dependence. When the influence of the full range of past trials was considered (SDE-all), the bias increased to about 40% (Figure 2A). These values were measured under filtering conditions that enhanced the expression of serial dependence (detailed in the next section, *Conditions enhancing SDE: influence of target visibility and location*). Importantly, these biases were not attributable to motor responses, as they persisted in a dual-task setup and were modulated by the spatial location and visibility of target stimuli, independent of motor actions (Figure 2).

### Conditions enhancing SDE: influence of target visibility and location

Biases were most pronounced under conditions where the current targets were barely visible (SOA < SOA_threshold_ + 20 ms) and prior targets were clearly visible (SOA > SOA_threshold_) with SDE-all reaching 40 ± 3% (Figure 2A; Supplementary Figure S2A). Biases were significantly reduced when the current targets were highly visible (SDE-all = 5 ± 1%; reduction: 35 ± 3%; t(49) = 11.9, p < 0.0001, Cohen’s d = 1.7) or when prior targets had low visibility (SDE-all = 5 ± 1%; reduction: 35 ± 3%; t(49) = 13.5, p < 0.0001, Cohen’s d = 1.9). In the 2loc condition, biases were predominantly location-selective, with significantly stronger effects when trial history originated from the same location (SDE-all = 41 ± 3%) compared to a diagonally opposite location (SDE-all = 15 ± 3%; reduction: 26 ± 3%; t(13) = 7.8, p < 0.0001, Cohen’s d = 2.1; Figure 2B; Supplementary Figure S2B).

### Decay of SDE over trials and with longer RT

Biases gradually decayed across successive trials but remained substantial, extending far into trial history (Figure 2 and Figure 6A). In the 1loc condition, biases were significant up to 4 trials back (p < 0.001 for N ≤ 4). In the 2loc condition, biases persisted longer, remaining significant up to 8 trials back (p < 0.001 for N ≤ 7; p < 0.05 for N = 8). For contra 2loc condition, biases were significant up to 5 trials back (p < 0.001 for N ≤ 2, p < 0.01 for N = 3, 4; p<0.05 for N=5). The dummy condition showed the most prolonged biases, with significant effects extending up to 9 trials back (p < 0.001 for N ≤ 6; p ≤ 0.01 for N = 7, 9), although the effect at 8-back was not significant (p = 0.77) and 10-back was borderline significant (p = 0.05). Our analysis focused on decay across trials rather than elapsed time, as doubling inter-trial interval had no impact on the short-history effects but only attenuated the long-history effects (as observed when comparing the two groups in the 1loc condition; see Supplementary Material Table S1 for group details).

Motivated by recently found mechanism-dependent relationship between response bias and reaction time (RT) (Dekel & Sagi, 2020), we examined the dependence of SDE on RT. We compared SDEs in trials with the fastest and slowest RTs. For each observer, trials were divided into quartiles based on RTs calculated separately for each training day, to account for overall reductions in RT with practice. SDEs were then computed using all trials from the fastest quartile (lowest 25%) and the slowest quartile (highest 25%) (Figure 3A). Recent SDEs were significantly stronger for fast RT (SDE-recent = 29 ± 2%) compared to slow RT (SDE-recent = 22 ± 2%), yielding a reduction of 7 ± 2% (t(49) = 3, p < 0.01, Cohen’s d = 0.4; Figure 3B, left panel; Supplementary Figure S2C), whereas distant SDEs did not significantly differ between RT conditions (SDE-distant = 9 ± 1% for both fast and slow RT; t(49) = 0.5, p = 0.64; Figure 3B, right panel; Supplementary Figure S2C).This difference between recent and distant SDEs suggests that they may arise from distinct underlying mechanisms. Observers from all conditions were combined in this analysis to increase statistical power, as separate analyses revealed a similar qualitative pattern across conditions: recent SDEs were stronger for fast compared to slow RTs, reaching significance in the 1loc and 2loc conditions but not in the dummy condition, likely due to smaller sample size. In contrast, distant SDEs showed no significant RT-related change in any condition.

**Figure 3.**
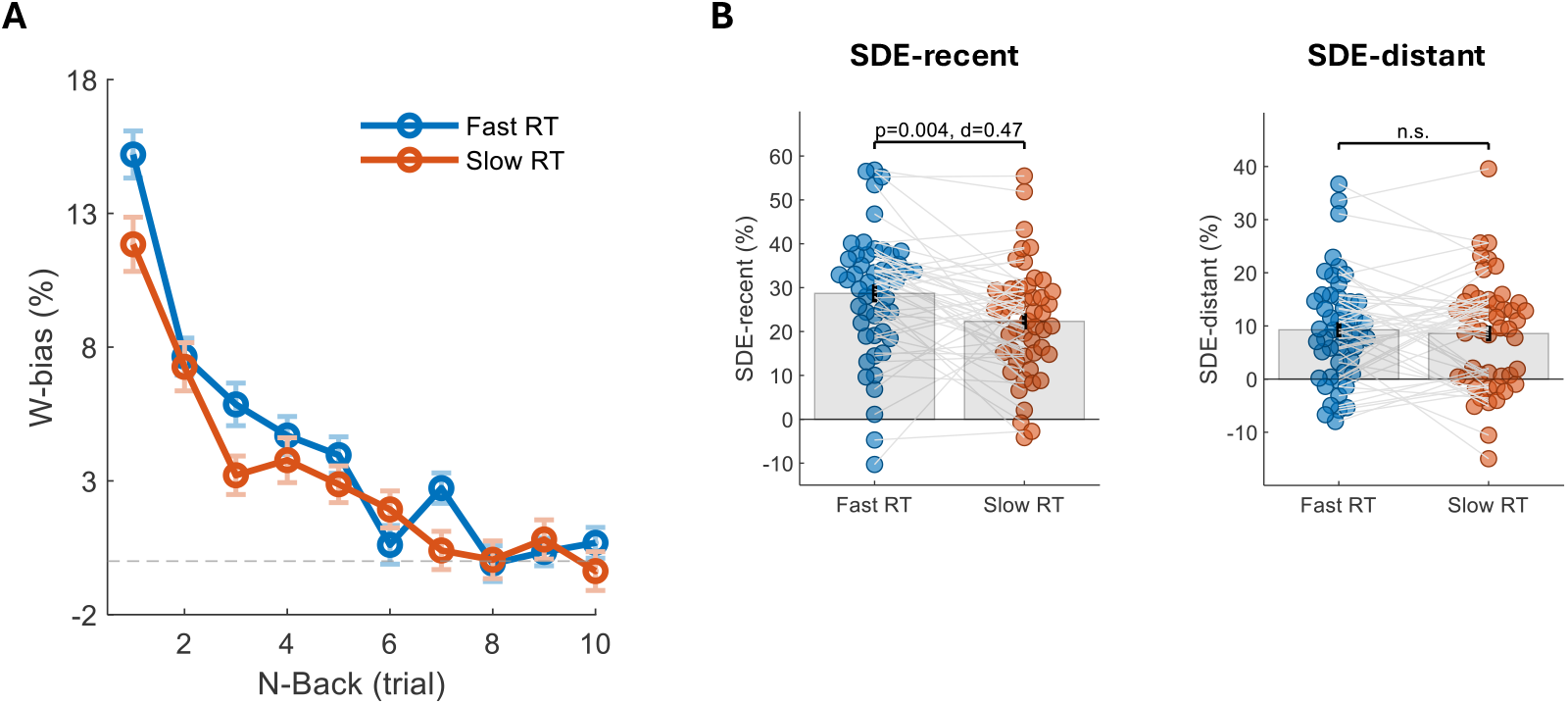
Effect of reaction time on SDEs. (A) SDEs (W-bias) as a function of N-back trial, calculated separately for the fastest (first 25%) and slowest (last 25%) RT quartiles, defined per day within each observer. SDEs were then computed on the corresponding fast and slow trials and averaged across observers. Biases were stronger for fast RTs, particularly at recent lags. (B) Paired comparisons of recent and distant SDEs for fast vs. slow RTs. Recent SDEs were significantly higher for fast RTs (left panel), whereas distant SDEs did not differ between RT conditions (right panel). Gray bars indicate group means; dots and connecting lines represent individual observers (*N = 50*).

**Figure 4.**
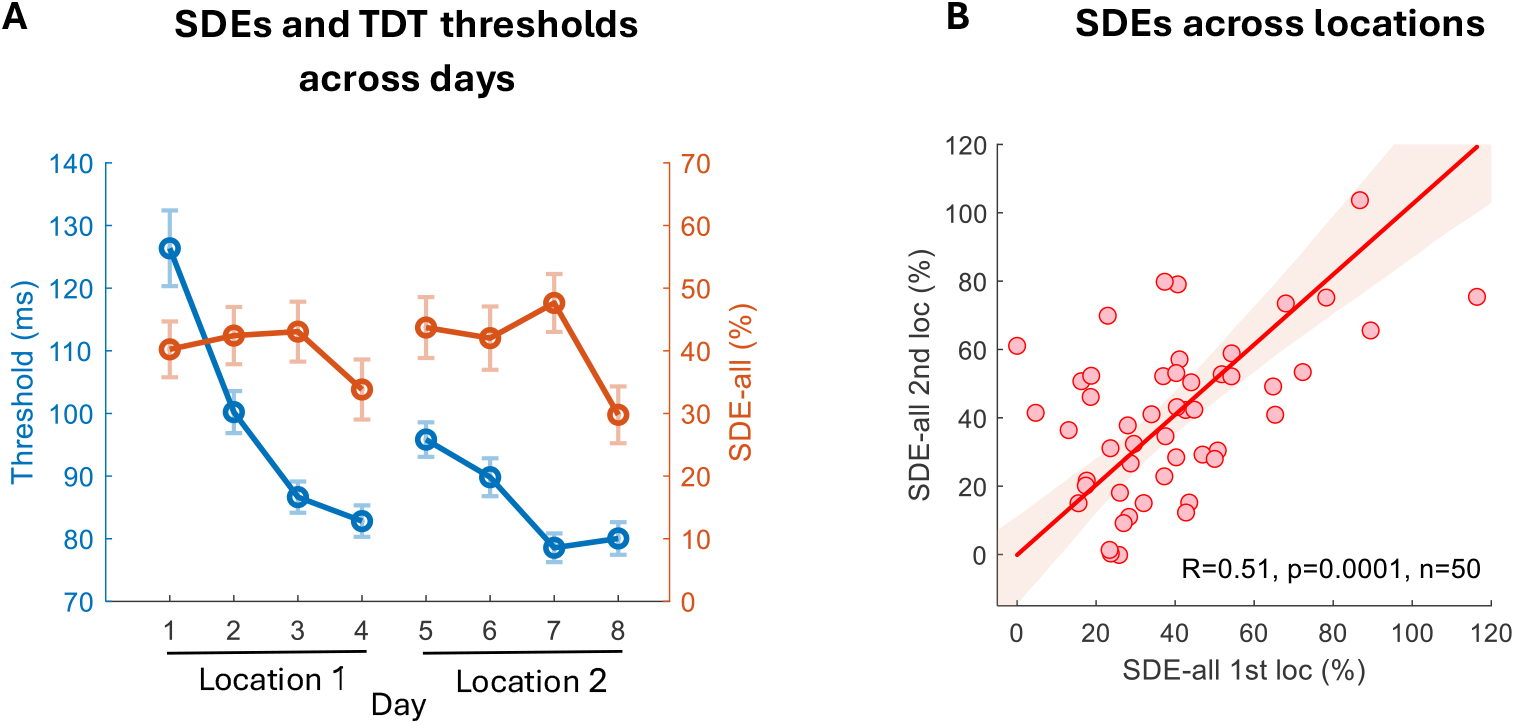
Dynamics of SDE across days and locations. (A) SDEs and TDT thresholds across days: Serial dependence (SDE-all, red) remained strong and consistent throughout the eight training days, despite large improvements in TDT thresholds (blue). A small reduction in SDE was observed across days and reached significance only between Days 7–8. The target location was changed after Day 4. (B) SDEs across locations: Correlation of SDE-all between the first (Days 1–4) and second (Days 5–8) trained locations across observers (*N = 50*). The strong correlation indicates that the magnitude of serial dependence is a stable observer-specific trait, consistent across retinotopic locations. In (B), shaded regions denote 95% bootstrap confidence sleeves around the orthogonal regression fit.

**Figure 5.**
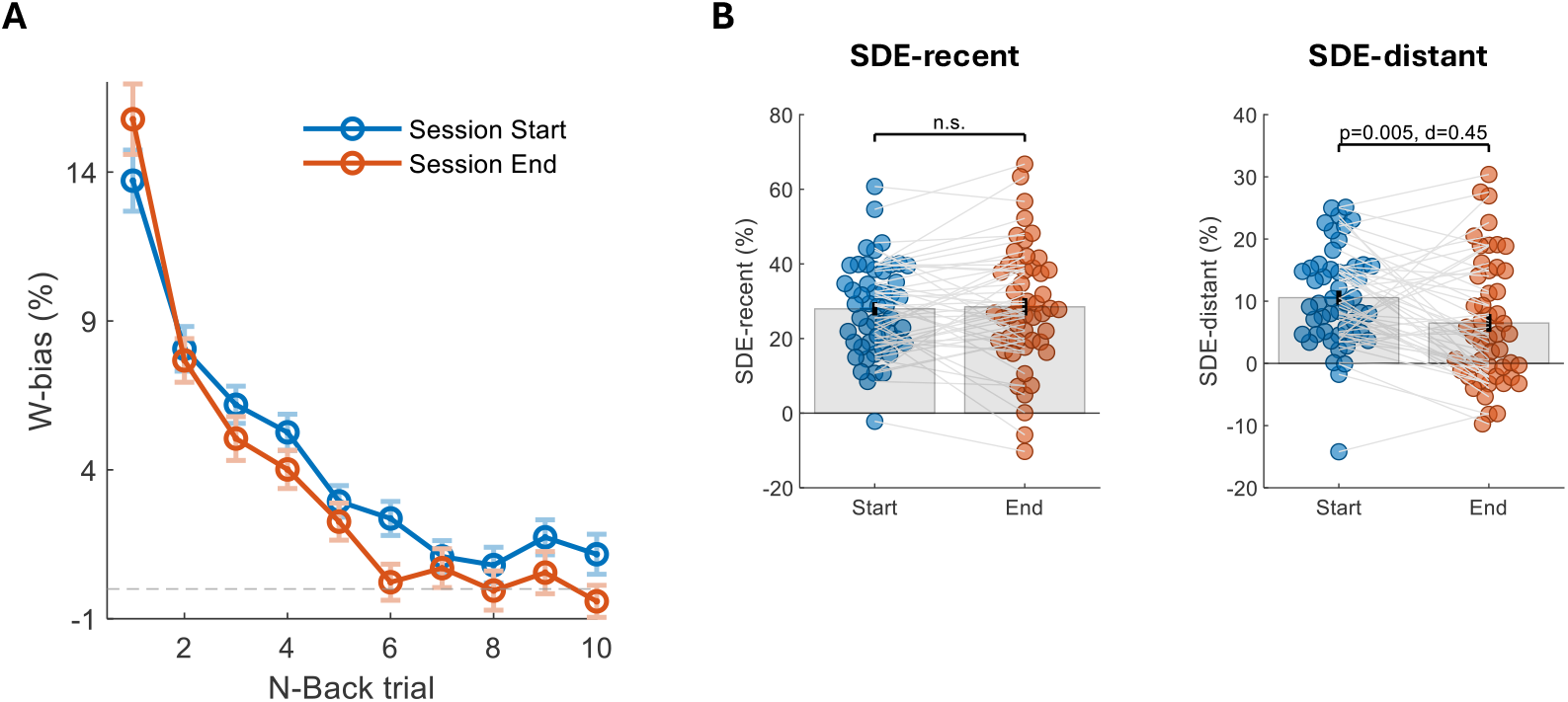
Within-session dynamics of SDE. (A) SDEs (W-bias) as a function of N-back trial, computed separately for the first (blue) and last (red) third of each session. Biases showed an overall decrease across the session, while the 1-back bias (W1) increased slightly. (B) Recent and distant SDE components. Recent SDEs remained stable across the session (left panel), whereas distant SDEs showed a significant reduction (right panel). This pattern is consistent with sensory adaptation developing over the course of the session, selectively attenuating serial dependence from more temporally distant trials (*N* = 50).

**Figure 6.**
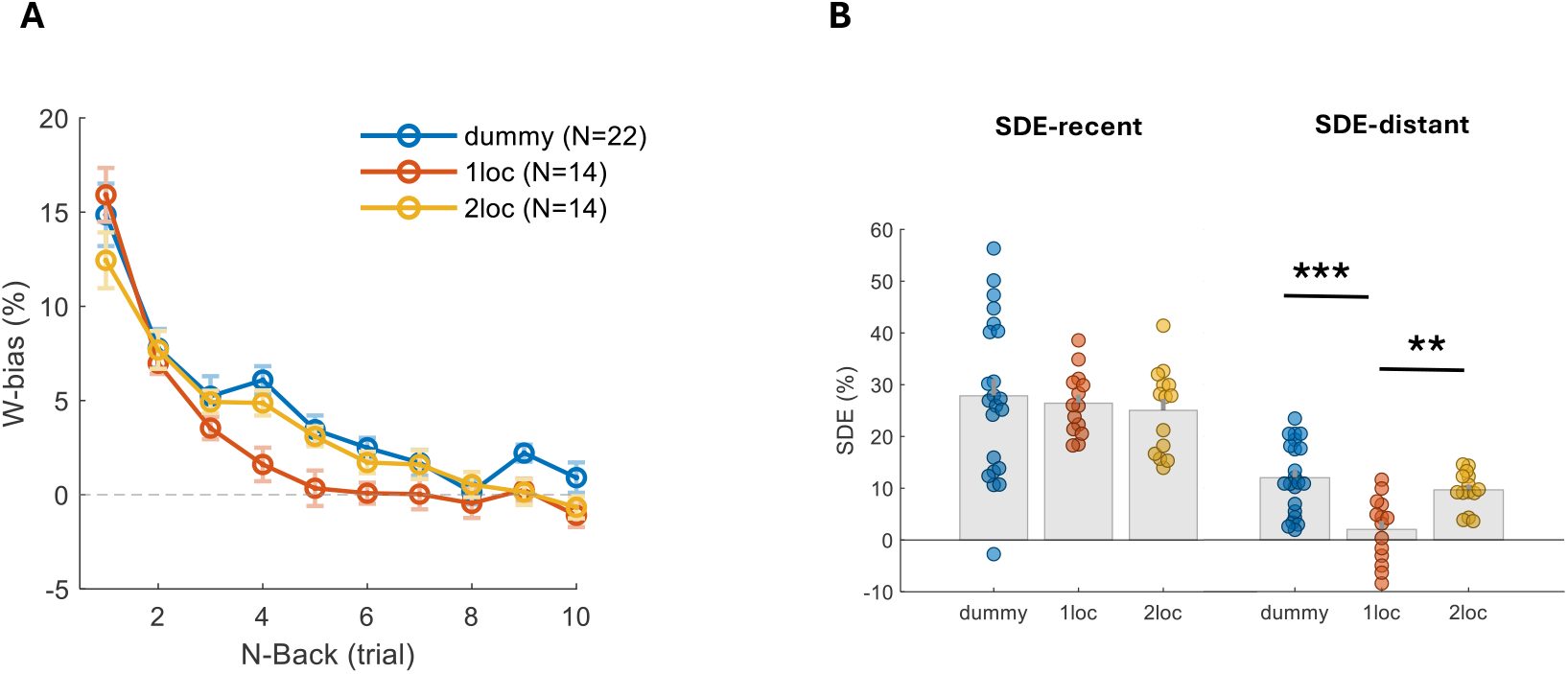
SDEs across experimental conditions and trial history. (A) Mean SDEs (W-bias) as a function of trial history (N-back) for the three experimental conditions: dummy (blue, *N* = 22), 1loc (red, *N* = 14), and 2loc (yellow, *N* = 14). In the 1loc condition, biases decayed more rapidly, while in the 2loc and dummy conditions they persisted further back in trial history. (B) SDEs across individual observers (*N = 50*), shown separately for recent lags (1–3 back; left panel) and distant lags (4–6 back; right panel). Each dot represents one observer; gray bars indicate group means. Recent SDEs were consistent across conditions, whereas distant SDEs were significantly stronger in the 2loc and dummy conditions compared to the 1loc condition (***p ≤ 0.001, **p ≤ 0.01).

**Figure 7.**
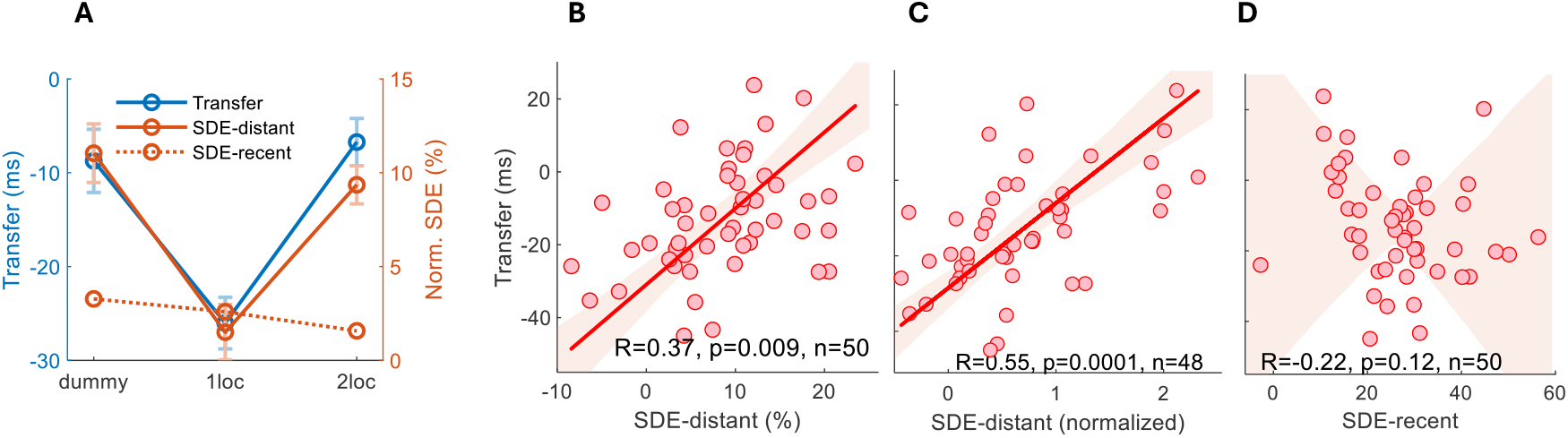
Relationship between SDE and learning transfer. (A) Group-level comparison of learning transfer (blue), SDE-distant (solid red), and SDE-recent (dotted red) across the three experimental conditions (dummy, 1loc, 2loc). Transfer and SDE values are plotted on separate axes, with SDE measures normalized by subtracting the mean of the 1loc condition. Conditions showing greater learning generalization (dummy, 2loc) also exhibited stronger SDE-distant effects. In contrast, SDE-recent was relatively constant across conditions, suggesting that generalization was primarily linked to distant serial dependence. (B) Across observers, learning transfer correlated positively with SDE-distant (r = 0.37, p < 0.01, *N = 50*), indicating that stronger distant serial dependence predicted greater generalization. (C) SDE-distant values were normalized to the 1-back effect to estimate the temporal decay constant of SDE, reflecting how long biases persisted across trials. Observers with longer decay constants showed greater learning transfer (r = 0.50, p < 0.001, *N = 48*; two outliers >10 SD excluded), indicating that extended temporal integration supports generalization. (D) No significant correlation was found between SDE-recent and learning transfer (r = –0.22, p = 0.12, *N = 50*), suggesting that recent serial dependence does not predict generalization. Learning transfer was defined as the change in TDT threshold between Day 4 (final day at the first location) and Day 5 (initial day at the second location), with negative values indicating performance loss. In B-D. shaded regions denote 95% bootstrap confidence sleeves around the orthogonal regression fit.

### Dynamics of SDE across days and locations

Despite randomized target orientations across trials, rendering past orientations irrelevant to current judgments, serial dependence biases remained strong and highly significant across all training days (p < 0.0001 for each day; Figure 4A). A two-factor repeated-measures ANOVA with Location (first vs. second retinotopic site) and Day-in-block (1–4) as within-subject factors revealed a significant main effect of Day-in-block, F(3, 147) = 4.7, p < 0.01, indicating modest decrease of SDE magnitude across training days. The main effect of Location was not significant (F(1, 49) = 0.08, p = 0.78), with comparable SDE-all values at the first trained location (Days 1–4: 40 ± 3%) and the second trained location (Days 5–8: 41 ± 3%). There was no Location × Day-in-block interaction (F(3, 147) = 0.47, p = 0.71), indicating similar temporal dynamics across locations. Post-hoc comparisons (Bonferroni-corrected) revealed a modest but statistically reliable decrease in SDE from Day 7 to Day 8 (the last two sessions at the second location; p < 0.01), whereas no other day-to-day differences reached significance. No significant correlation was found between biases and SOA thresholds across observers (r = -0.13, p = 0.37, average across days 1-8), nor between biases and improvements in performance at the first location (r = -0.09, p = 0.54, average across days 1-4), suggesting that the magnitude of serial dependence does not predict the overall amount of perceptual learning (Supplementary Figure S1).

Across the eight days of training, SDEs thus remained robust even as texture-discrimination thresholds improved markedly from Day 1 (126 ± 6 ms) to Day 8 (80 ± 2 ms; F(7, 343) = 41.77, p < 0.0001; Figure 4A). Biases were highly correlated across locations (r = 0.51, p < 0.001; Figure 4B), suggesting that the magnitude of serial dependence reflects a stable observer-specific trait consistent across retinotopic locations.

### Within-session SDE dynamics

Within-session analyses (averaged across all training days and conditions) showed that serial dependence biases remained significant throughout sessions but decreased by approximately 17% from the beginning to the end. To track such bias changes, while having a sufficient number of trials for bias analysis, each session was divided into three parts. Biases in the first third of trials (SDE-all = 43 ± 3%) were significantly higher than in the final third (SDE-all = 36 ± 4%), yielding an 8 ± 3% reduction (t(49) = 2.4, p < 0.05, Cohen’s d = 0.3; Figure 5A). This decrease may reflect sensory adaptation developing over the course of the session, diminishing serial dependence. The reduction was selective to distant lag history: SDE-distant decreased significantly from 11 ± 1% to 7 ± 1% (t(49) = 2.9, p < 0.01, Cohen’s d = 0.4; Figure 5B, right; Supplementary Figure S2D), whereas SDE-recent remained stable (28 ± 2% in both segments; t(49) = 0.31, p = 0.76; Figure 5B, left). Notably, the 1-back bias (W1) showed a slight increase from start to end (2 ± 1%; t(49) = 2.2, p < 0.05, Cohen’s d = 0.3; Figure 5A), indicating that within-session adaptation primarily attenuates the influence of distant SDEs while leaving immediate history effects intact, or even slightly enhanced.

Observers from all conditions were combined in this analysis to increase statistical power, as separate condition-level analyses revealed the same trend (no change in SDE-recent and >30% reduction in SDE-distant), but these did not reach significance, likely due to smaller sample sizes.

### SDE differences between conditions and learning generalization

A comparison across the three experimental conditions revealed similar magnitudes of SDE-recent (dummy: 28 ± 3%; 1loc: 26 ± 2%; 2loc: 25 ± 2%; F(2,47) = 0.26, p = 0.77; Figure 6B left). In contrast, SDE-distant differed significantly between conditions (dummy: 12 ± 2%; 2loc: 10 ± 1%; 1loc: 2 ± 2%; F(2,47) = 12.47, p < 0.001; Figure 6B right). Post-hoc Tukey tests confirmed that SDE-distant was significantly lower in the 1loc condition compared to both the dummy (p < 0.001) and 2loc (p < 0.01) conditions, likely due to stronger sensory adaptation caused by repeated stimulation at a fixed location in the 1loc setup. In the letter discrimination (fixation control) task, which involved identical foveal stimuli across all conditions, no significant differences were observed for either SDE-recent (dummy: 6 ± 2%; 1loc: 8 ± 2%; 2loc: 8 ± 2%; F(2,47) = 0.40, p = 0.674) or SDE-distant (dummy: 2 ± 1%; 1loc: 2 ± 1%; 2loc: 3 ± 1%; F(2,47) = 0.59, p = 0.56). The smaller biases in this task likely resulted from higher overall performance levels and ceiling effects.

We next examined whether the greater learning generalization observed in the 2loc and dummy conditions (Harris et al., 2012) is linked to the stronger distant serial dependence found in those conditions (Figure 7A).

Supporting this, we found that learning transfer correlated positively with SDE-distant across observers (r = 0.37, p < 0.01; Figure 7B) and with SDE-distant values normalized to the 1-back effect to estimate the temporal decay constant of SDE (r = 0.50, p < 0.001; Figure 7C). In contrast, SDE-recent showed no positive correlation (r = -0.22, p = 0.12; Figure 7D), and became significantly negative when one outlier (>3 SD) was excluded (r = -0.32, p < 0.05, N = 49), suggesting that recent-trial biases, being more closely tied to the current stimulus, may have a weaker and less consistent relationship with learning generalization (Figure 7A).

## Discussion

Our investigation of serial dependence in the texture discrimination task (TDT) reveals robust perceptual biases toward the orientation of previously presented targets, extending up to 10 trials back under certain conditions. Notably, these biases persisted despite randomized target orientations and significant improvements in performance across training days, suggesting that serial dependence is a fundamental feature of visual processing, largely unaffected by task demands or learning. While our findings are based on the texture discrimination task, we expect the link between long-range serial dependence and learning generalization to extend across perceptual domains. Serial dependence and perceptual learning have been documented for numerous features including orientation, numerosity, face identity, and auditory pitch (Manassi et al., 2023; Lau & Maus, 2019; Sagi, 2011), suggesting that future work could further test this link as our framework predicts. Considering the universality of learning mechanisms in the brain (Censor et al., 2012), we suggest that this newly established link is not limited to visual perception but rather a general property of human behavior.

While previous studies typically report a 3-back limit for serial dependence (Fischer & Whitney, 2014; John-Saaltink et al., 2016; Lau & Maus, 2019; Manassi et al., 2019, 2023), our results reveal exceptionally long memory traces, extending up to 8-back in the 2loc condition and up to 9-back in the dummy condition. Consistent with earlier work, we found that the reliability of both current and prior target stimuli affected bias magnitude. Biases increased when current targets were less visible (Ceylan et al., 2021; Cicchini et al., 2017, 2018; Manassi et al., 2018) and when prior targets were more visible (Pascucci et al., 2019; Van Bergen & Jehee, 2019). This pattern aligns with Bayesian models of perception (Kersten et al., 2004; Knill & Pouget, 2004) which propose that perceptual estimates under uncertainty integrate current sensory evidence with prior information. Additionally, we observed spatial selectivity in the 2loc condition, with stronger biases when prior and current targets appeared at the same location, in agreement with previous reports (Collins, 2019; Fischer & Whitney, 2014; Fornaciai & Park, 2018; John-Saaltink et al., 2016; Manassi et al., 2019). Most interestingly, in our experiments without feedback on the texture task, the experimental conditions yielding the strongest bias were also reported to enhance learning in the absence of feedback (Liu et al., 2012).

Importantly, our findings reveal a functional link between serial dependence and perceptual learning. Serial dependence extends beyond transient perceptual biases and influences long-term learning, specifically, the extent to which learning generalizes to new, untrained locations. Observers trained under conditions that promote generalization (‘2loc’, ‘dummy’; Harris et al., 2012) exhibited significantly stronger and more temporally extended serial dependence from distant trial histories (4–6 back). Conversely, consistent stimulus repetition in the ‘1loc’ condition, which promotes location-specific learning, was associated with a shorter temporal span of serial dependence, likely due to the stronger sensory adaptation (Censor & Sagi, 2009; Harris et al., 2012; Ofen et al., 2007). Across individuals in all conditions, greater distant SDEs predicted greater learning transfer. These results suggest a unified mechanism in which short-term memory traces, as reflected in serial dependence, can either accumulate to support generalization or be truncated, possibly by adaptation, limiting learning to the trained context. Limited generalization is often attributed to smaller or less variable training sets in machine learning (Ying, 2019), and in perceptual learning (Sagi, 2011), which can lead to overfitting. A similar principle may apply here: the shorter integration window in 1loc limits the accumulation of informative variability, promoting overfitting and thus reducing generalization, thus a longer history adds little value (Tavoni et al., 2022).

To our knowledge, no previous study has experimentally linked serial dependence to long-term perceptual learning. However, a theoretical framework proposed by Pascucci et al. (2019) connects short-term history biases to learning mechanisms, suggesting that what appears as bias in serial dependence tasks may actually reflect the process by which the visual system updates its decision templates, the same mechanism thought to underlie perceptual learning (Dosher & Lu, 1998; Kuai et al., 2013). Specifically, they argue that serial dependence arises from the reinstatement of previously informative sensory channels, effectively reusing feature weights that were beneficial in previous trials. Similarly, Talluri et al. (2018) found that observers selectively overweight evidence consistent with prior choices, and Murai & Whitney (2021), using classification-image analysis, demonstrated that serial dependence reshapes the perceptual templates applied to upcoming stimuli. Supporting this view, Urai et al. (2019) fitted bounded-accumulation models and showed that choice history is best explained by a history-dependent change in evidence accumulation (implemented as a drift bias), rather than merely shifting the starting point of the decision process (criterion bias). This result is in agreement with our RT analysis showing that distant SDEs are RT-independent, a marker of drift bias (Dekel & Sagi, 2020). In addition, the recent trials seem to introduce criterion shifts (starting point bias in the drift diffusion model), indicated by the larger biases found for fast RTs. Together, these findings suggest that serial dependence directly alters how sensory information is weighted and interpreted. We suggest that these updated decision templates subserve perceptual learning.

The persistence of serial dependence during eight days of training with random stimulus sequences, where it does not contribute to online performance, but rather increases decision noise, suggests that the assumption of environment stability is hardwired into the brain, or that these biases may serve a broader function beyond optimizing immediate performance. Consistent with our approach, recent models reframe serial dependence as a memory-driven phenomenon, not as an optimal inference about the external world (Barbosa & Compte, 2020), but as a consequence of internal mechanisms shaped by how recent perceptual states are encoded and maintained over time (Kalm & Norris, 2018).

Our findings reveal a functional/mechanistic dissociation between short- and long-range serial dependence. Only recent SDEs were modulated by reaction time, presenting stronger biases with faster responses, suggesting that these biases are due to shifts in decision criteria (Dekel & Sagi, 2020). In contrast, distant SDEs were found to be RT independent, suggesting that these biases are a result of neuronal reweighting (Dekel & Sagi, 2020). Importantly, only distant SDEs predicted learning transfer, while recent SDEs remained stable across conditions and were unrelated to generalization. The within session dynamics showed distant SDEs, but not recent SDE, to decline with training, thus effectively reducing the SDE range, consistent with the increased learning specificity observed in perceptual learning with extensive learning (Sagi, 2011). This pattern suggests a functional distinction: recent biases may relate more to prior stimulus statistics, whereas temporally extended biases may support the integration of sensory evidence required for efficient perceptual learning. Previous studies also point to distinct timescales in serial dependence. For example, Lieder et al. (2019) showed that perceptual biases reflect processes operating over different timescales that vary across clinical populations: individuals with ASD rely less on recent trials but show intact long-term integration, whereas individuals with dyslexia exhibit the opposite pattern. Thus, in the absence of adaptation, we expect learning in ASD to generalize, as indeed was recently found (Harris et al., 2015). Fritsche et al. (2020) proposed a model in which perceptual history influences current biases through both short-term Bayesian decoding and longer-term efficient encoding, aligning with our observed dissociation between recent and distant SDEs. While these converging findings support distinct mechanisms for recent and distant SDEs, our correlational approach cannot definitively establish causality, and targeted experimental manipulations would further strengthen these interpretations.

Our findings offer a new insight into the mechanisms of perceptual learning. While traditional theories explain learning specificity through local changes at the site of target encoding (Karni & Sagi, 1991), the formation of location-specific decision templates (Dosher & Lu, 1999), or both (Karni & Sagi, 1993; Watanabe & Sasaki, 2015), we propose a unified mechanism. Specifically, we suggest an account based on a single decision template that learns the discrimination task by classifying neuronal response features as signaling vertical or horizontal texture targets. These templates generalize across retinal locations of equal eccentricity but not across locations with different eccentricities (Harris & Sagi, 2018). However, when trained with targets in a fixed location, the decision template may become overfitted to features that are specific to that location, limiting generalization (Sagi, 2011). For learning to generalize, multiple samples (trials) must be integrated over time to filter out local noise. Our results show that decision biases are integrated linearly over trials, suggesting efficient temporal integration over many trials in conditions that support learning generalization. In contrast, reduced integration, due to adaptation, or increased inhibition, may produce classifiers that rely on location-specific noise (Mollon & Danilova, 1996). We suggest that previous reports of learning generalization can be explained by a modulation of temporal integration. This includes short training phases that are stopped before adaptation takes over, showing generalization to other retinal locations (Censor & Sagi, 2009; Karni & Sagi, 1993), short pre-training phases enabling generalization across visual tasks (Zhang et al., 2010) and other paradigms that effectively reduce sensory adaptation (reviewed in Sagi, 2011) and by that allow serial dependence to accumulate. Our findings thus provide empirical support for a unified mechanism that governs both specific and generalized learning through modulation of temporal integration. We further speculate that the integration window is affected by the balance between excitation and inhibition (E/I balance) in the visual cortex, shown to affect learning stabilization in TDT (Shibata et al., 2017; Tamaki et al., 2020). Computational models implementing trial-by-trial reweighting (e.g, Petrov et al., 2005) with adaptation dependent reweighting, can potentially account for SDE decay profiles and their relationship to generalization, providing quantitative predictions for future experiments (or for already existing experimental data reanalyzed for SDE). Weights update dynamics may affect network flexibility and generalization. To account for the present results within this general framework, we assume that consistent stimulus repetition (triggering sensory adaptation) stabilizes learning by lowering the gain of network update. Reduced sensory adaptation (disinhibition) allows for increased plasticity producing network dynamics allowing for faster adjustment to new stimuli. This update mechanism is expected to introduce serial dependencies with a temporal scale defined by E/I balance. To test the plausibility of this approach, we constructed a simple computational model of learning presented in Supplementary results. This toy model, based on modelling learning in volatile environments (Piray and Daw, 2020), predicts serial dependence produced by template update, with a magnitude correlated with generalization. An important property of this model is the continuous plasticity, that is the system does not stop updating its templates. Attractive serial dependence emerges as an immediate consequence of ongoing template learning: informative (high SOA, easy) trials selectively update orientation templates, and subsequent ambiguous (low SOA, noisy) trials read out these updated templates, producing an attractive bias without requiring explicit feedback or top-down control. In addition, we may consider biases due to updated priors, as commonly assumed in the SDE literature. This positions serial dependence as a signature of the same flexibility mechanisms that support adaptive learning in nonstationary environments.

In summary, we show that long-range serial dependence predicts learning transfer, supporting the view that short-term memory contributes directly to long-term learning. By connecting serial dependence with learning, our findings bridge a key theoretical gap and suggest that the integration of past experience plays a crucial role in determining the specificity or generalization of learning.

## Supplementary material: Tables and Figures

**Table S1.** Group assignments in each condition.

| Condition | Group | N | Comments |
| --- | --- | --- | --- |
| 1loc | 1loc-standard | 7 |  |
|  | 1loc-dark | 7 | Double inter-trial period |
| 2loc | 2loc-standard | 7 |  |
|  | 2loc-long | 7 | Twice the number of trials |
| Dummy | dummy-standard | 7 | Background of horizontal bars |
|  | dummy-90reBck | 5 | Background of vertical bars |
|  | dummy-90reTarget | 5 | Background of diagonal bars (angled at 45°) |
|  | dummy-local | 5 | Local background with only 5 × 5 texture elements |

**Figure S1.**
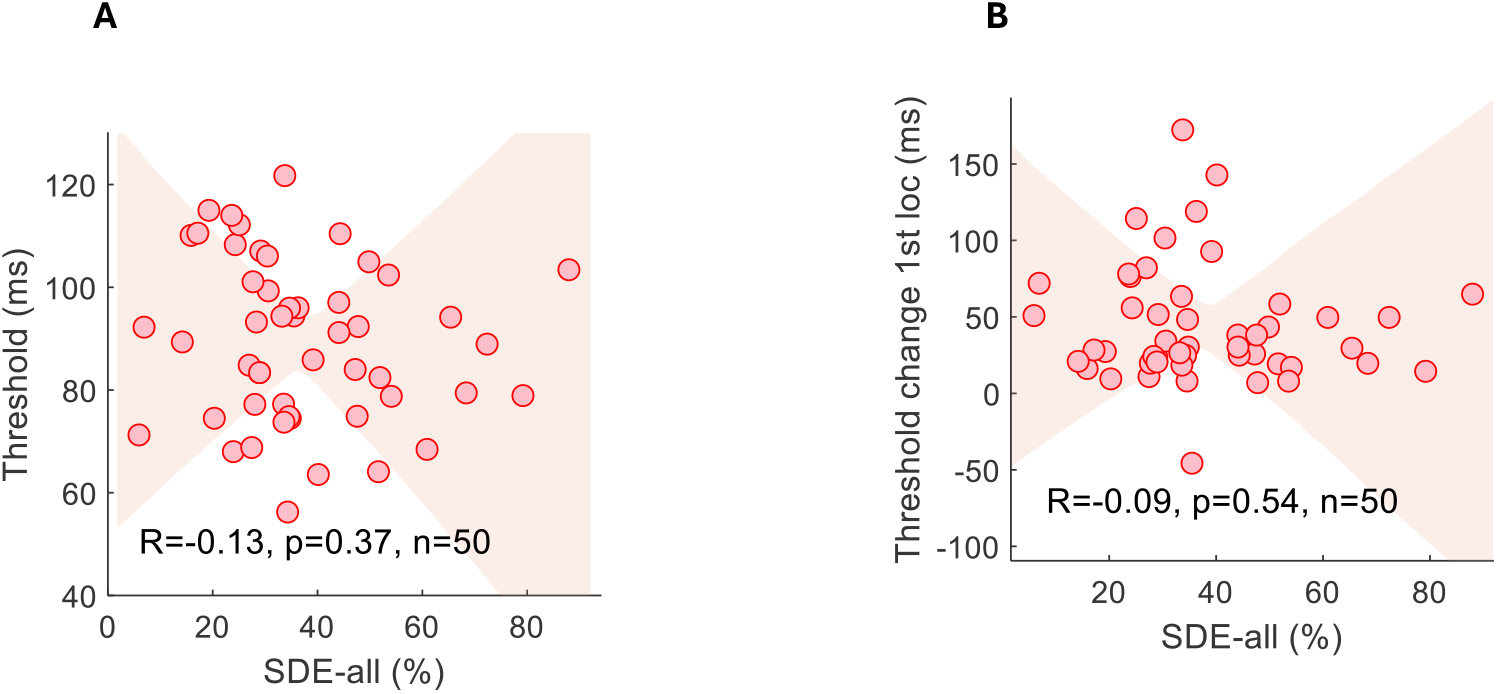
Relationship between TDT Threshold and SDEs. (A) Correlation between average TDT thresholds (Days 1–8) and SDE-all across observers (n = 50). No significant relationship was found (r = –0.13, p = 0.37). (B) Correlation between threshold improvements at the first location (Day 1 to day 4) and SDE-all. No significant relationship was found (r = –0.09, p = 0.54), suggesting that the magnitude of serial dependence does not predict the overall amount of perceptual learning.

**Figure S2.**
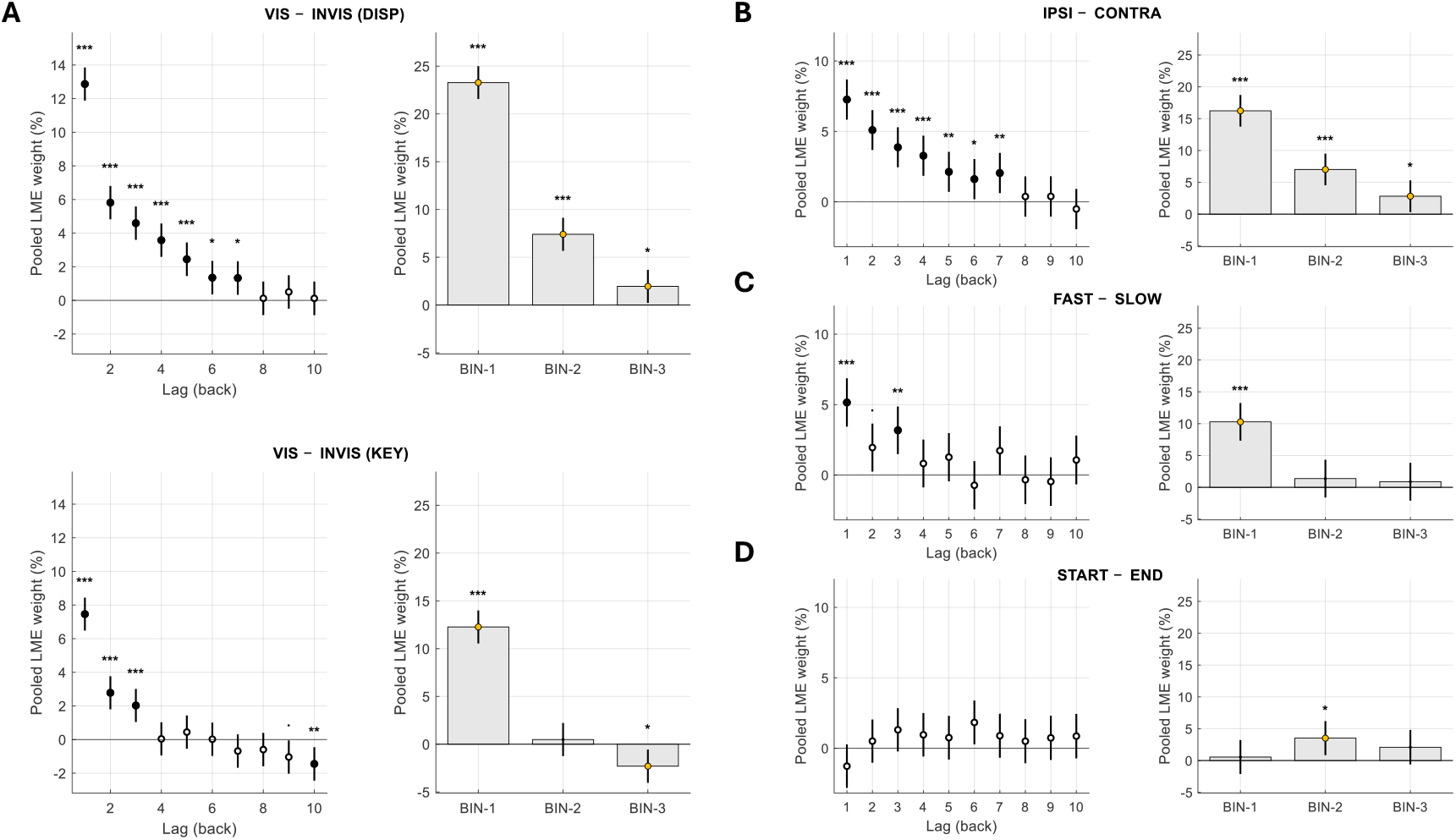
Serial-dependence contrasts from pooled LME estimates. Each subfigure shows SDE across individual lags (left) and lag bins (right). For each contrast, lag-specific history weights from the linear mixed-effects (LME) models were computed as the difference between conditions over the ten preceding trials (lags 1–10). Differences were computed either within the same model (VIS − INVIS, IPSI − CONTRA) or between independent model fits trained on matched subsets of trials (FAST − SLOW, START − END). Estimates were pooled across observers using inverse-variance weighting (grand-pooled across groups for VIS − INVIS; cross-fit and pooled across observers for FAST − SLOW and START − END; IPSI − CONTRA computed for loc2). Values are pooled contrast estimates (Δ ± SE, % bias). Significance is based on FDR-corrected *q*-values (Benjamini–Hochberg across lags). Bin panels summarize BIN1 (1–3 back), BIN2 (4–6 back), BIN3 (7–9 back). **(A) VIS – INVIS**. SDEs were reduced after low-visibility trials, with a larger reduction for display-based (target) than for key-based (response) histories; key-driven SDEs showed a small significant increase at distant lags for invisible trials. **(B) IPSI − CONTRA**. SDEs were stronger for ipsilateral than contralateral locations, indicating spatial specificity. **(C) FAST – SLOW**. Trials were split into fastest and slowest reaction-time quartiles per observer and day. Biases were stronger for fast trials, particularly at recent lags (SDE-recent). **(D) START – END**. SDEs computed for the first vs. last third of each session showed an overall decline toward session end, with slight increase in the immediate (1-back) bias. The reduction reached significance only for BIN2 (SDE-distant). Error bars denote ±1 SEM across observers. Asterisks indicate FDR-corrected significance (***q < .001, **q < .01, *q < .05).

**Figure S3.**
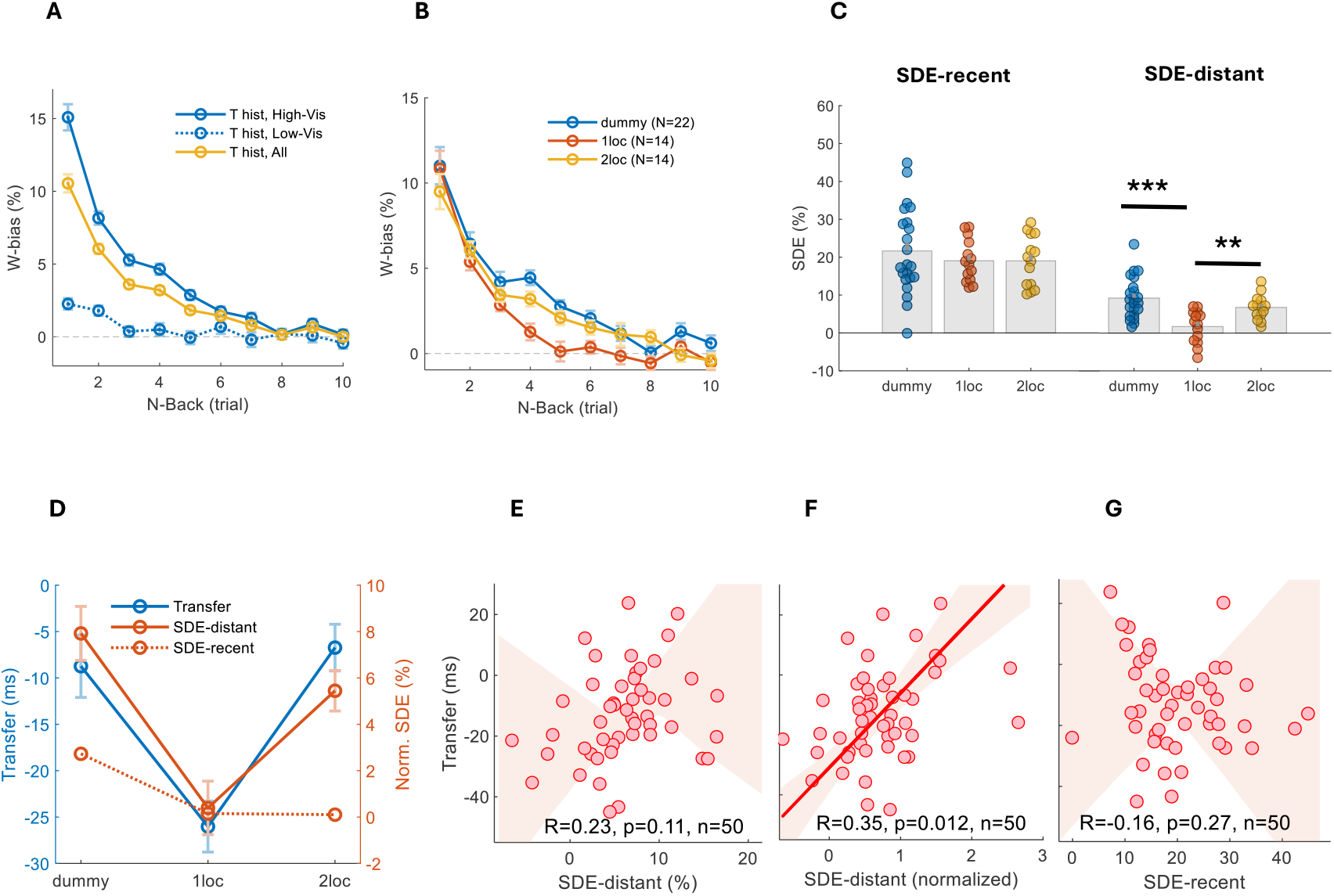
Serial dependence effects using all prior-trial history (unfiltered). **(A)** W-bias as a function of n-back lag, comparing high-visibility (solid blue), low-visibility (dashed blue), and all prior trials (yellow). Data pooled across conditions. **(B)** W-bias decay across n-back lags for each experimental condition (dummy, 1loc, 2loc) using all prior trials. As with the high-visibility prior analysis, biases decayed more rapidly in the 1loc condition, while persisting further back in trial history for the 2loc and dummy conditions. **(C)** Individual SDE values by condition for SDE-recent (left) and SDE-distant (right). Bars indicate group means; dots represent individual observers. As with the high-visibility prior analysis, recent SDEs were consistent across conditions (dummy: 29 ± 3%; 1loc: 29 ± 2%; 2loc: 28 ± 2%; F(2,47) = 0.08, p = 0.92), whereas distant SDEs were significantly stronger in the 2loc and dummy conditions compared to the 1loc condition (dummy: 12 ± 2%; 1loc: 3 ± 1%; 2loc: 11 ± 1%; F(2,47) = 11.27, p = 0.001). Post-hoc Tukey tests confirmed that SDE-distant was significantly lower in the 1loc condition compared to both the dummy (***p < 0.001) and 2loc (**p < 0.01) conditions. **(D)** Group-level comparison of learning transfer (blue), SDE-distant (solid red), and SDE-recent (dotted red) across the three experimental conditions (dummy, 1loc, 2loc). Transfer and SDE values are plotted on separate axes, with SDE measures normalized by subtracting the mean of the 1loc condition. Conditions showing greater learning generalization (dummy, 2loc) also exhibited stronger SDE-distant effects. In contrast, SDE-recent was relatively constant across conditions, suggesting that generalization was primarily linked to distant serial dependence. **(E)** Across observers, learning transfer was not significantly correlated with SDE-distant (r=0.23, p=0.11, N=50), suggesting that using all prior-trial history reduces the effect. **(F)** SDE-distant values were normalized to the 1-back effect to estimate the temporal decay constant of SDE, reflecting how long biases persisted across trials. Observers with longer decay constants showed greater learning transfer (r=0.35, p=0.012, N=50), similar to the high-visibility prior analysis. **(G)** No significant correlation was found between SDE-recent and learning transfer (r=-0.16, p=0.27, N=50), suggesting that recent serial dependence does not predict generalization. In E-G, shaded regions denote 95% bootstrap confidence sleeves around the orthogonal regression fit.

## Supplementary material: modelling

Here we describe a simple model of learning that generates persistent serial dependence (SDE) as a consequence of weight update. There is no intention of fitting the experimental data but rather showing that SDE can arise as a consequence of weights update in a learning model. We here describe the learning mechanism, followed by model equations and simulations.

In the experiments simulated here, on each trial observers are visually presented with one of two stimulus categories, Vertical or Horizontal. Targets are masked by noise of variable impact, determined by Stimulus Onset Asynchrony (SOA) between the target and mask stimuli. On each trial, observers decide whether the presented stimulus is Vertical or Horizontal. There was no response feedback in the experiments and in the simulations. The main results to account for are the long range SDE and the correlation between SDE and learning transfer.

In the simulations, observers are assumed to maintain two online category templates, *μ*_*A*_ and *μ*_*B*_, updated from trial-by-trial sensory samples via responsibility-weighted, Kalman-like template updates. Updates follow a variant of the Volatile Kalman Filter (VKF; Piray & Daw, 2020). In our simulations the categories are orientations: the observer keeps two running estimates of what Vertical (category A) and Horizontal (category B) targets “look like” along a 1-D decision axis (orientation evidence in our experiments). These internal templates correspond to current estimates of the category means, *μ*_*A*_ and *μ*_*B*_.

On each trial, the visual system generates an internal sensory response *x*_*t*_, added with stimulus-independent sensory noise σ. Using the two templates and the current category prior, a posterior responsibility can be computed

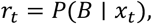

i.e., how strongly the current sample supports category B rather than A. This responsibility is used for:

- Decision: report B if *r*_*t*_ > 0.5.
- Learning: update both templates, but with responsibility-dependent weights. If *r*_*t*_ ≈ 1, the B-template is updated strongly and the A-template minimally; if *r*_*t*_ ≈ 0.5, both updates are small; if *r*_*t*_ ≈ 0, the A-template is updated strongly and the B-template minimally.

There are easy and hard trials defined by SOA: easy trials (high SOA, the noisy mask does not interfere with the target stimulus) tend to generate samples *x*_*t*_ farther from the decision boundary, yielding more extreme responsibilities (*r*_*t*_ near 0 or 1), whereas hard trials (low SOA, target corrupted by noise) yield samples closer to the decision boundary with *r*_*t*_ near 0.5. As a result, the internal state (templates, and therefore the implied boundary between them) shows jumps followed by persistence: informative/easy samples produce larger template shifts, while ambiguous/hard samples produce little change.

Serial dependence follows directly. An easy trial shifts the internal boundary; subsequent hard trials are ambiguous and therefore sensitive to the exact positioning of the boundary. Because hard trials do not strongly update the templates, the boundary persists and the bias extends across multiple lags. In the simulations this link is verified by computing the boundary shift

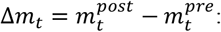

and showing that Δ*m*_*t*_, differs after EASY-A versus EASY-B trials. Moreover, for EASY→HARD pairs, Δ*m*_*t*−1_ predicts the probability of reporting B on the next hard trial.

Of interest here is that the same mechanism that generates SDE also determines flexibility (generalization). In VKF-like updates, volatility enters as a process-noise term *Q*_*t*_ that controls the Kalman gains (see below ‘Volatility’). When mismatch accumulates (operationalized below as a large prediction error 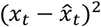; e.g., after a rotation/location switch), *Q*_*t*_ increases, gains increase, and the templates move faster toward the new mapping—yielding rapid relearning, considered here as learning generalization if very fast (e.g. ‘dummy’ condition). When volatility is depressed (e.g. ‘1loc’ condition), gains are smaller and the template state is more conservative: boundary shifts produced by informative events are smaller (weaker SDE, bidirectional), and re-learning after a change is slower. *The positive SDE observed with high volatility is a result of the high update gain that tends to overshoot the updated template behind its expected ‘true’ value. Thus, SDE and re-learning are two observable consequences of the same underlying state-update machinery—one read out as a bias on ambiguous trials, the other read out as the speed of convergence after a mapping change*.

In addition, we have a sticky prior over categories. Instead of keeping the prior p_B_ fixed at 0.5, the observer maintains an online estimate of how likely category B is: after each trial, the prior shifts slightly toward the current posterior r_t_ and then relaxes back toward 0.5. This, as described in the SDE literature, provides a source of serial dependence that biases inference directly (via prior odds), even in the absence of template shifts.

## Technical description

### State variables

- Template means: μ_A,t_, μ_B,t_
- Template uncertainties: P_A,t_, P_B,t_
- Category prior: p_B,t_
- Volatility state: v_t_

### Trial order

At trial t: (1) observe *x*_*t*_; (2) compute r_t_ using the pre-update templates and prior; (3) decide A or B; (4) update volatility; (5) update templates; (6) update sticky prior (if enabled).

### Decision (uses pre-update templates)

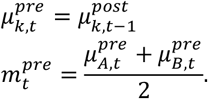

### Inference

Likelihoods use fixed internal σ:

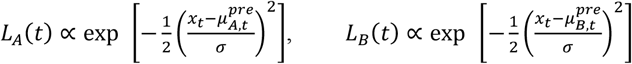

Posterior responsibility:

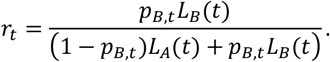

Decision: report B if r_t_ > 0.5.

#### Confidence gate

To reduce learning from ambiguous stimuli, we optionally apply a confidence gate based on posterior distance from indifference:

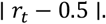

A bounded confidence gate is:

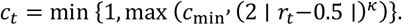

This downweights learning from ambiguous (dummy) trials (*r*_*t*_ ≈ 0.5).

### Volatility (flexibility)

Prediction:

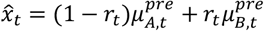

Volatility update (*v*_*t*_ increases when new observation is far from the predicted, 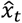):

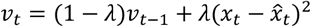

Process noise:

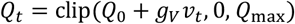

### Template updating

Prediction uncertainty (updated by process noise):

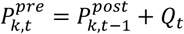

Observation uncertainty (fixed):

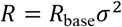

Responsibility-weighted effective noise:

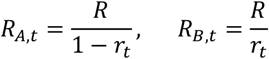

Kalman gains:

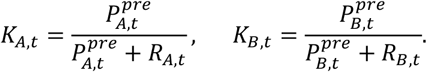

Updates:

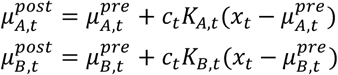

Boundary step:

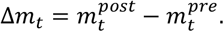

Large mismatch → larger *v*_*t*_→ larger *Q*_*t*_→ faster re-learning.

### Sticky prior dynamics

The prior is updated as a leaky average of recent posteriors:

History integration:

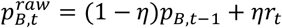

Relaxation toward neutrality:

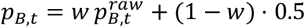

Where

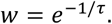

So:

*η* controls how strongly the current posterior influences the next prior.

*τ* controls how slowly the prior relaxes back toward 0.5.

This prior is used in the next trial’s responsibility calculation.

### Serial dependence mechanisms

Serial dependence in this model can arise from two coupled mechanisms:

A. Template-shift mechanism. Easy trials induce nonzero boundary steps Δ*m*_*t*_, while hard trials do not strongly overwrite the boundary. Because 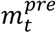 is autocorrelated, conditioning on a past easy A vs B trial predicts current boundary position, biasing ambiguous decisions. Volatility *Q*_*t*_ contributes to serial dependence because it sets the Kalman gains: higher *Q*_*t*_ increases the amplitude of easy-trial boundary steps (stronger bias).
B. Sticky-prior mechanism. The prior *p*_*B,t*_ integrates recent posteriors and directly biases the next trial’s inference via prior odds:

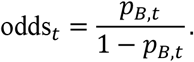

Even if templates were frozen, this would generate serial dependence. When both mechanisms are active, SDE reflects both persistent boundary shifts and prior drift.

### Simulation results

There was no attempt to fit the data, but rather to demonstrate the similarity between model behavior and that observed in the experiments. The main interest here is in showing the ability of the model to imitate the experimental results from the ‘1loc’ and the ‘dummy’ conditions, when controlling volatility (*Q*_0_, *g*_*V*_). Simulations followed the scheme presented above. There were 100 experiments of 100K trials with easy, hard and noisy/dummy (mean=0) trials.

**Figure S4:**
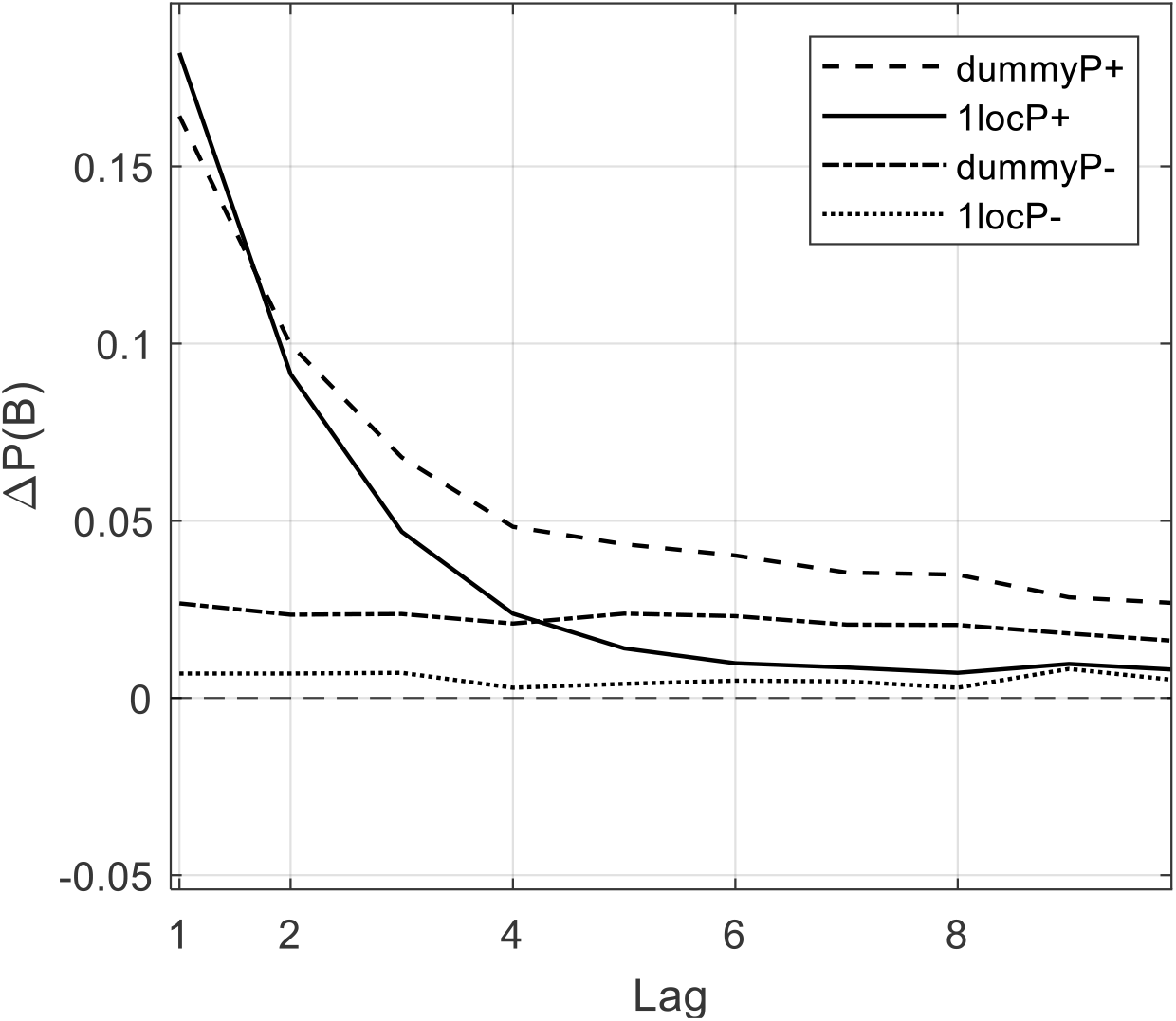
Simulated bias as a function of lag (trials) for four conditions: (1) high volatility with sticky prior, simulating the dummy condition, (2) low volatility with sticky prior, simulating the 1loc condition, and (3) high volatility without sticky prior, and (4) low volatility without sticky prior. Condition 3 (dummyP-) reflects pure template-learning SDE (i.e., serial dependence arising from template updates alone). Results averaged across 100 runs (different random seed), 100K trials each. *g*_*V*_ = 0.005 for dummy, *g*_*V*_ = 0 for 1loc. Compare with Figure 6A in main text.

**Figure S5:**
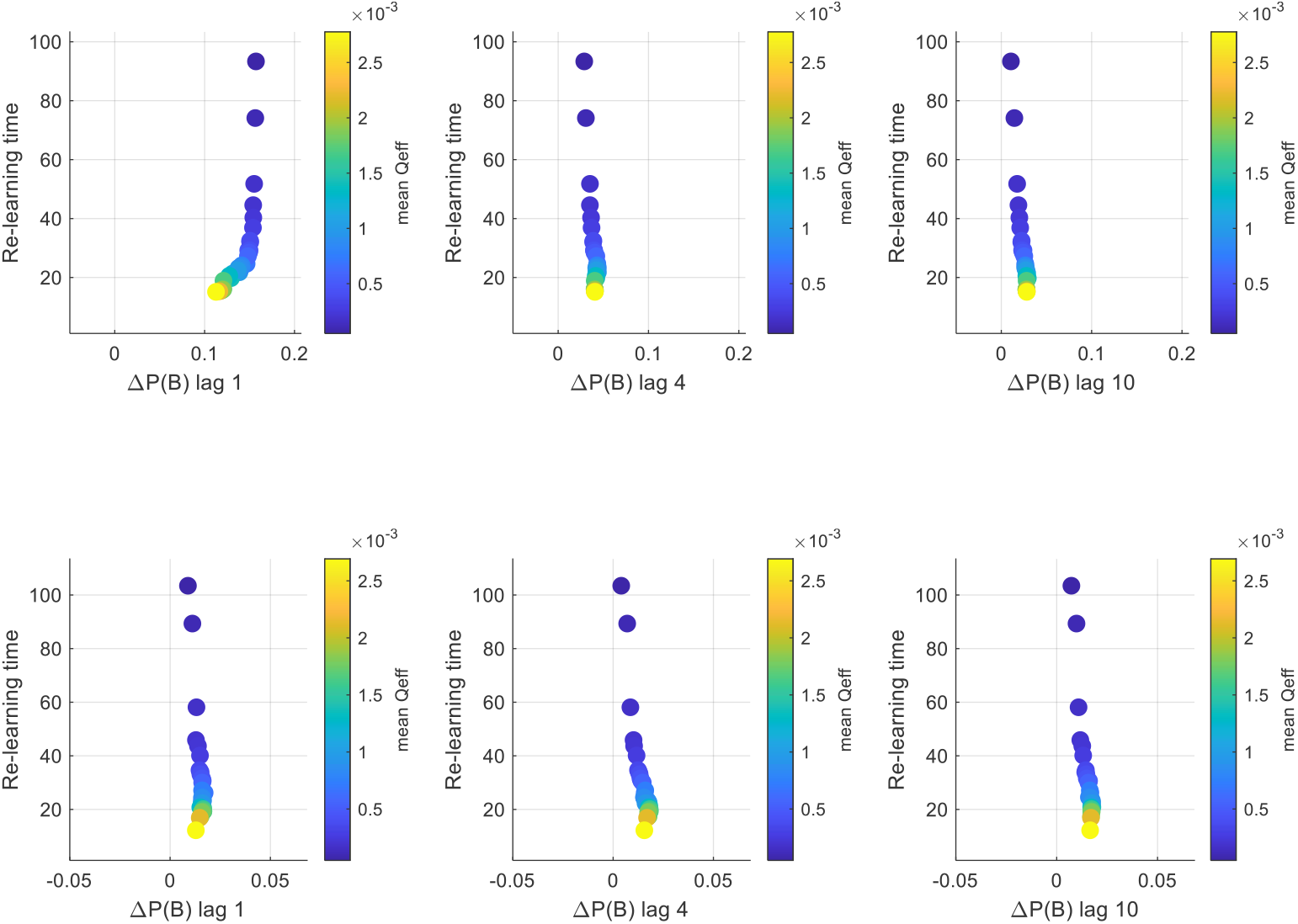
SDE and learning generalization. Plotted are re-learning times (in trial units) as a function of bias, for three lags (1, 4, 10), with (top) and without (bottom) sticky priors. The results qualitatively demonstrate the phenomenon observed in the experiments: larger biases correlate with faster re-learning upon stimulus change. Color code represents mean volatility (Q) corresponding to specific relearning-lag pairs.

## Methods

### Overview

We simulated a nonstationary 1D categorization task with online template learning and quantified lagged sequential dependence effects (SDE) on current **hard A/B** trials near the true category boundary. Simulations were Monte Carlo over independent random seeds and additionally included a third **noise (dummy)** trial type embedded in the stream.

### Environment and trial generation

Each simulation comprised *T* = 100,000 trials with regime switches governed by a Bernoulli hazard *H* = 1/10000 per trial. Regime 1 had true category means (*μ*_*A*_, *μ*_*B*_) = (−1, +1); regime 2 had (*μ*_*A*_, *μ*_*B*_) = (0, +2). On each trial, with probability *π*_*N*_ = 0.25 the sample was a **noise** trial generated as 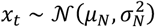 with *μ*_*N*_ = 0 and *σ*_*N*_ = 0.1; otherwise the trial was an A/B trial with category B drawn with base rate *π*_*B*_ = 0.5 and sample generated as 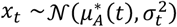 or 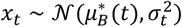. A reliability schedule controlled *σ*_*t*_ for A/B trials: a warm-up period of *t*_Warm_ = 100 trials used “easy” trials only *σ*_Easy_ = 0.5; thereafter each A/B trial was easy with probability *p*_Easy_ = 0.25 and hard otherwise, using *σ*_Hard_ = 1.0. Easy/hard tags were defined for A/B trials only (noise trials excluded). Noise trials were part of the temporal stream and counted toward lags; in the default configuration they updated the learner. Noise trials simulated dummy trials and were not present when the standard 1loc condition was simulated.

### Distance-to-boundary binning

For each trial we computed the true midpoint boundary 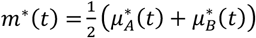 and distance *d*_*t*_ =| *x*_*t*_ − *m*^∗^(*t*) |. Trials were binned into **near/mid/far** using thresholds thr_1_ = 0.5 and thr_2_ = 1.0 (near if *d*_*t*_ ≤ thr_1_, mid if thr_1_ < *d*_*t*_ ≤ thr_2_, far if *d*_*t*_ > thr_2_). Primary SDE analyses focused on *current* hard A/B trials in the near bin.

### Learner (online template model)

The observer maintained estimates of A and B template means 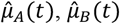, scalar uncertainties *P*_*A*_(*t*), *P*_*B*_(*t*), and (optionally) a dynamic prior *p*_*B*_(*t*). On each trial it computed a soft responsibility *r*_*t*_ = *P*(*B* | *x*_*t*_) from Gaussian-shaped likelihoods centered on the current templates with fixed sensory scale *σ*_sense_ =0.6 and prior *p*_*B*_, and generated a binary response B when (*r*_*t*_>0.5), or A otherwise. Template updates were Kalman-like uncertainties were inflated by a process-noise term *Q*_*t*_, responsibility-weighted effective observation noises were formed, Kalman gains computed, and both templates were updated proportional to prediction errors. Update magnitude was modulated by a confidence gate *c*_*t*_ derived from | *r*_*t*_ − 0.5 | (parameters confMin=0.05, confPower=2). Flexibility was controlled by a VKF-like volatility estimate: a latent volatility state *v*_*t*_ tracked an exponential moving average of squared prediction errors with forgetting l=0.05, and process noise was set to *Q*_*t*_ = clip(*Q*_0_ + *g*_*V*_*v*_*t*_, [0, *Q*_max_]) with *Q*_max_ = 0.2. Baseline observation noise was R_base_=0.1 (scaled by 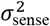). Learning occurred on all trials, including noise. A sticky-prior module (disabled for template-shift-only analyses) updated *p*_*B*_ toward *r*_*t*_with etaPrior=0.5 and relaxed it toward 0.5 with time constant tauPrior=3. Initial learner state was 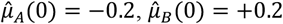, *P*_*A*_(0) = *P*_*B*_(0) = 1.0, *p*_*B*_(0) = 0.5, and *v*_0_ = 0.

### Monte Carlo and analysis windows

We simulated *n*_Seeds_ = 100 independent environments and ran the learner on each. All analyses used a burn-in period *t*_Burn_ = *t*_Warm_ + 1000 = 1100 trials to exclude initialization transients. Lag analyses used maximum lag *L*_max_ = 10 and required at least minCount=200 samples per condition.

### Sequential dependence metric

For each lag *ℓ*, we computed the bias in B reports on a set of *current* trials conditioned on the true label of the trial *ℓ* steps back, restricted to cases where that previous trial was **easy A/B**. Specifically, we estimated *b*_*B*_(*ℓ*) = 2*P*(report B | current mask, prev easy true *B*) − 1 and *b*_*A*_(*ℓ*) = 2*P*(report B | current mask, prev easy true *A*) − 1, and combined them as 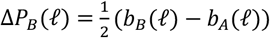. This analysis was carried out on current **hard A/B** trials in the **near** bin, with SEM computed across seeds (decisions on noisy, dummy, trials were excluded).

### Mechanistic boundary-step coupling

To link SDE to learning-driven boundary motion, we defined the learned midpoint 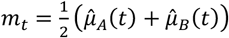 and boundary step Δ*m*_*t*_ = *m*_*t*_ − *m*_*t*−1_. We measured Δ*m*_*t*_ on easy A/B trials split by true label, and quantified coupling by binning Δ*m*_*t*−1_ from previous easy trials and computing *P*(report B at *t*) on the immediately subsequent hard A/B trial.

### Adaptation time after regime switches

Flexibility was quantified as relearning time following regime switches using the template error 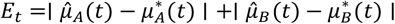. Switches were separated by at least 5000 trials. For each switch, baseline error was the mean error in the pre-window; relearning time was the first post-switch time when error returned to within tolFrac=0.15 of baseline, averaged across switches and seeds.

### Parameter sweep

To characterize the bias–adaptation tradeoff, we ran a grid sweep over volatility parameters *Q*_0_ ∈ {0,5 ⋅ 10^−5^, 10^−4^, 2 ⋅ 10^−4^, 5 ⋅ 10^−4^, 10^−3^} and *g*_*V*_ ∈ {0,5 ⋅ 10^−4^, 10^−3^, 2 ⋅ 10^−3^, 4 ⋅ 10^−3^}, recomputing mean adaptation time and SDE at lags {1, 4, 10} (default near-bin hard A/B) for each parameter pair; points were visualized as adaptation time versus SDE magnitude, color-coded by mean effective process noise ⟨*Q*_*t*_⟩.

### Compact parameter list (defaults)

Monte Carlo: seedBase 42, *N* = 100. Environment: *T* = 100,000, *t*_warm_ = 100, hazard *H* = 1/10000, *π*_*B*_ = 0.5, regimes (*μ*_*A*_, *μ*_*B*_) = (−1, +1) and (0, +2). Reliability: *p*_easy_ = 0.25, *σ*_easy_ = 0.5, *σ*_hard_ = 1.0. Noise trials: *π*_*N*_ = 0.25, *μ*_*N*_ = 0, *σ*_*N*_ = 0.1. Analysis: *t*_burn_ = 1100, *L*_max_ = 10, minCount=200, bins thr_1_ = 0.5, thr_2_ = 1.0. Learner: sigSense=0.6, R_base=0.1, confidence on (confMin=0.05, confPower=2), VKF on, lambda=0.05, Q_0_=10^−5^, Q_max_=0.2, initial 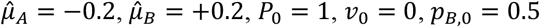. Sticky prior for optional runs: etaPrior=0.5, tauPrior=3 (disabled for template-shift-only analyses). *g*_*V*_ = 0.005 for dummy, *g*_*V*_ = 0 for 1loc.

